# WDR81 represses IKK mediated expression of pro-survival genes to regulate apoptosis

**DOI:** 10.1101/2025.03.27.645662

**Authors:** Sannoong Hu, Pranav Danthi

## Abstract

Apoptosis is a common host response to virus infection. The extent and timing of apoptosis following infection is controlled by the balance between the strength of signals that activate death signaling and those that promote the survival of cells. In many cell types, infection with Mammalian orthoreovirus (reovirus) results in induction of cell death by apoptosis late in infection. In this study, we uncovered that WD repeat-containing protein 81 (WDR81) is required for apoptosis induction after reovirus infection. The requirement for WDR81 for apoptosis induction is not unique to reovirus because cells lacking WDR81 are also resistant to apoptosis induced by other agonists that trigger the extrinsic apoptotic pathway. We find that in cells deficient in WDR81, expression of several pro-survival genes is upregulated. The expression of these genes is controlled by the inhibitor of kB kinase complex (IKK)-Nuclear Factor of kB (NFκB) signaling pathway. When IKK signaling is blocked in WDR81-deficient cells, pro-survival gene expression is restored to normal levels and the cells regain their susceptibility to reovirus-induced death receptor-triggered apoptosis. Our work uncovers a new function for WDR81 in controlling apoptosis. Additionally, it reveals a previously unknown link between endosomally localized protein WDR81 and IKK-NFκB signaling.

**Author Summary:** Virus infection often results in death of the infected cells. Cell death prior to generation of virus progeny limits the spread of infection to neighboring cells and therefore can be beneficial to the host. However, cell death might also cause tissue destruction and could contribute to viral disease. It is therefore important to understand how cell death is controlled. Here, we uncover a cell death regulating role for WDR81 – a cellular protein that has not been previously implicated in affecting cell death. We find that when this protein is absent, cells express a much greater level of survival signals. These survival signals prevent efficient induction of cell death. By investigating how these survival signals are expressed, we reveal a new link between WDR81 and NFκB, a well-known cellular survival pathway.

## Introduction

Apoptosis is a well-studied form of programmed cell death. Apoptosis is characterized by activation of a cascade of proteases called caspases^1,2^. The activation of these caspases ultimately leads to the death of the cell in a non-inflammatory manner^2^. The apoptosis pathway can be triggered by intrinsic or extrinsic pathways. Extrinsic signals such as tumor necrosis factor (TNF) can bind to their receptors and trigger the activation of an initiator caspase, caspase-8. In its activated form, caspase-8 cleaves effector caspases, caspase-3 and caspase-7. The intrinsic pathway involves the mitochondria and results in the activation of caspase-9, which, in turn, also activates effector caspases. Activated effector caspases cleave numerous cellular components ultimately leading to cell death^2^.

Cells express proteins that prevent the spontaneous activation of caspases^2–4^. The expression of these pro-survival gene products is usually mediated by the IKK complex via the NFκB transcription factor^3,4^. Thus, in the absence of a cell death trigger, the cell continues to grow and divide without activating caspases. Apoptosis can be triggered to overcome survival signals by extrinsic signals like tumor necrosis factor (TNF) or TNF related apoptosis inducing ligand (TRAIL) or by intrinsic imbalances such as stress or intracellular pathogens like viruses^5–7^. Virus infection and replication within a cell is often detected by PAMPs (Pathogen Associated Molecular Patterns) which trigger a variety of cellular immune responses including cell death^5,8^. Mammalian orthoreovirus (reovirus) infection triggers apoptosis^9–11^. While reovirus activated apoptosis is not associated with limiting viral replication within the infected cell, it has been associated with disease pathogenesis in mouse models^12–14^. Reovirus induced apoptosis requires activation of both the extrinsic and the intrinsic apoptotic pathways^11,15–17^. While a few viral and cellular factors are implicated in this process^10,18–22^, the exact manner in which apoptotic pathways are activated in reovirus infected cells is not understood.

We previously identified a function for a host protein, WDR81, in reovirus infection^23^. WDR81 regulates endosomal maturation^24^. It is present in the phosphoinositide-3-kinase (PI3K) complex on the outer leaflet of early endosomes and inhibits the function of Class III PI3Ks, thus allowing the maturation of the early endosome to late endosome^24^. In the absence of WDR81, reovirus virions attach to cell surface receptors, reach the endosome and are disassembled to generate entry intermediates called infectious subviral particles (ISVPs) by the action of endosomal proteases^23^. However, the ISVPs are trapped in the endosomes and fail to enter the cytoplasm and launch infection. When infection is launched with in vitro generated ISVPs, which do not require endosomal uptake, infection proceeds normally^23^. These data implicate WDR81 in regulating productive trafficking of reovirus particles through the endosomal pathway during the early stages of reovirus infection.

In this study, we analyzed the role of WDR81 in reovirus induced programmed cell death. We demonstrate that even though WDR81 is dispensable for infection by reovirus ISVPs, it is required for ISVP-induced apoptosis. We show that in the absence of WDR81, other extrinsic death agonists also fail to trigger apoptosis. Our data shows that the absence of WDR81 results in increased expression of pro-survival genes. We demonstrate that blocking the IKK-NFκB pathway normalizes the expression of these pro-survival genes and restores the capacity of extrinsic death triggers to induce apoptosis. Together, our study describes a novel function of WDR81 in apoptosis. Further, it reveals a previously unknown connection between WDR81 and the IKK-NFκB signaling pathway.

## Results

### WDR81 is required for reovirus induced programmed cell death

WDR81, a host protein critical for early-to-late endosome maturation, plays a key role in cellular trafficking^24^. Previous work demonstrated that WDR81 is essential for infection by reovirus virions but dispensable for infection by Infectious Subviral Particles (ISVPs), an entry intermediate generated via in vitro chymotrypsin digestion of virions^23^. To confirm this phenotype, control cells (WT MEFs) and cells deficient in WDR81 (ΔWDR81 MEFs) were infected with reovirus virions or ISVPs. Virus infectivity was measured 24 h post infection by indirect immunofluorescence. Consistent with previous work^23^, reovirus virions showed lower infectivity in ΔWDR81 MEFs compared to WT MEFs while infection with reovirus ISVPs showed no difference in infectivity between WT and ΔWDR81 MEFs (Fig. 1A). Virus production was also quantified by measuring viral titer 24 h post infection by plaque assay. Consistent with the previous results and indirect immunofluorescence, in comparison to WT MEFs, reovirus virions produced ∼0.7 log_10_ lower infectious virus in the ΔWDR81 MEFs while infection with reovirus ISVPs showed no difference in the amount of infectious titer produced in both cell types (Fig. 1B). To study the role of WDR81 in reovirus induced programmed cell death, WT MEFs and ΔWDR81 MEFs were infected with reovirus virions or ISVPs and incubated for 48 h. Permeability of host cell membranes to Sytox Green nucleic acid dye, which only stains nuclei of dead or dying cells^25^ was evaluated as a measure of cell death. This later time point was chosen as MEFs infected with reovirus do not show detectable cell death at 24 h post infection (data not shown). When infected with virions, in comparison to WT MEFs, ΔWDR81 MEFs showed significantly fewer dead cells (Fig. 1C). This result was expected as reovirus virions are unable to establish infection or replicate in ΔWDR81 MEFs (Fig. 1A and 1B). Interestingly, upon infection with reovirus ISVPs, ΔWDR81 MEFs still showed a significantly lower number of dead cells in comparison to the WT MEFs (Fig. 1C). This result was unexpected since reovirus ISVPs are capable of successful replication in ΔWDR81 MEFs, as shown in Fig. 1A and 1B. These results suggest that WDR81 plays a role in reovirus induced programmed cell death. Because ISVPs efficiently infected ΔWDR81 MEFs, for the remainder of this study, we used ISVPs as a tool to dissect the role of WDR81 in cell death induction.

**Figure 1.**
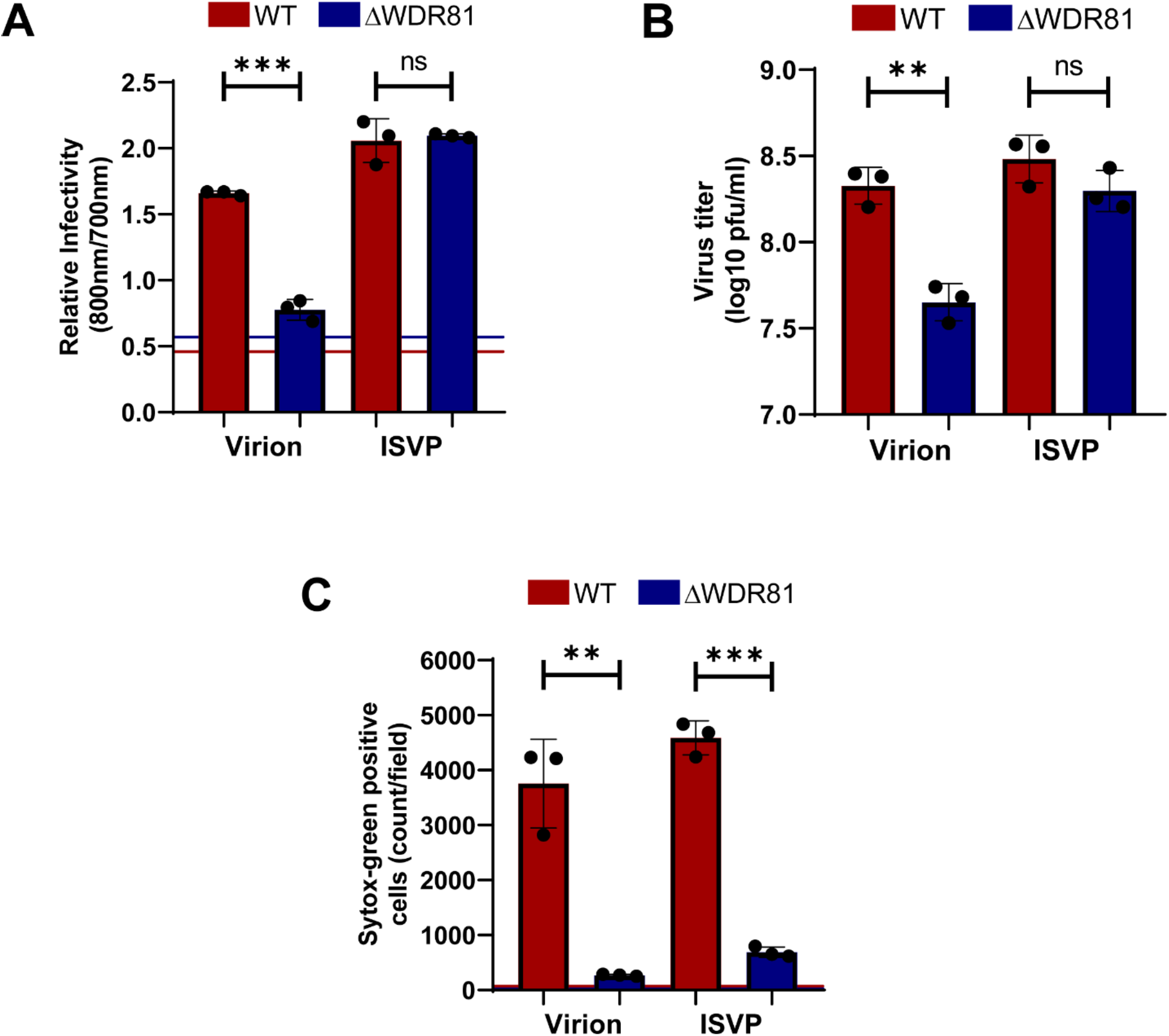
WDR81 is required for ISVP induced cell death. (A) MEFs were adsorbed with virions or ISVPs of reovirus T3D^CD^ at MOI 10 PFU/cell. Cells were then incubated for 24 h at 37°C. Infectivity was measured in fixed cells by indirect immunofluorescence using LI-COR Odyssey Scanner. Relative infectivity was calculated using the ratio of signal at 800nm (reovirus) to signal at 700nm (cells). Horizontal lines represent the ratio for mock infected cells for each cell type. Error bars represent SD. ***, P<0.0005 by student t-test. (B) MEFs were adsorbed with virions or ISVPs of reovirus T3D^CD^ at MOI 10 PFU/cell. Cells were incubated at 37°C for 24 h. Virus titer was measured by plaque assay on L929 cells. Error bars represent SD. **, P<0.005. (C) MEFs were adsorbed with virions or ISVPs of reovirus T3D^CD^ at MOI 10 PFU/cell. Cells were incubated for 48 h in media supplemented with Sytox– green (Sytox G) (50 nM). Cell death was quantified 48 h post infection by analyzing the number of Sytox G positive cells per field. Horizontal lines represent signals for mock infected cells for each cell type. Error bars represent SD. ***, P<0.0005 and **, P<0.005 by student t-test.

### WDR81 is required for reovirus induced caspase-3/7 activity

Reovirus induces cell death either through apoptosis, involving both the intrinsic and the extrinsic pathway, or necroptosis, mediated by RIPK3 and MLKL activation^9,26^. Reovirus infection of MEFs produces morphological and biochemical changes in cells that resemble apoptosis^22^. To determine whether reovirus infection induces apoptosis in MEFs, we assessed the effect of ZVAD-fmk^27^, a broad-spectrum caspase inhibitor, on cell death 48 h post-infection. As shown in Fig. 2B, ZVAD-fmk significantly suppressed ISVP-induced cell death in MEFs without affecting infection efficiency (Fig. 2A). These data suggest that MEFs undergo cell death by apoptosis following infection with reovirus ISVPs. Because ΔWDR81 cells fail to undergo cell death following infection by ISVPs, our results suggest that WDR81 is required for efficient induction of apoptosis. A hallmark of apoptosis is the activation of effector caspases – caspase-3 and caspase-7. To determine whether effector caspases are activated, WT and ΔWDR81 MEFs were infected with ISVPs. The activity of effector caspases in cell lysates at 48 h post infection was measured using a caspase-3/7 substrate which shows enhanced luminescence upon cleavage. Reovirus ISVP infection of WT MEFs resulted in a ∼2.5-fold increase in caspase-3/7 activity, compared to only a ∼1.5-fold increase in ΔWDR81 MEFs (Fig. 2C). Caspase-3/7 activation was also quantified using a cell-permeable substrate that fluoresces upon cleavage by active caspases. Upon infection of WT MEFs with reovirus ISVPs, we detected a large number of cells positive for caspase-3/7 activity. In contrast, ΔWDR81 MEFs exhibited markedly fewer cells with active caspase-3/7 (Fig. 2D). These data suggested that WDR81 is required for ISVP induced caspase-3/7 activation. To confirm the effectiveness of ZVAD-fmk, we also assessed caspase activation by ISVPs in the presence of ZVAD-fmk. ZVAD-fmk was efficiently able to inhibit caspase-3/7 activation (Fig. 2C and D). Together these data suggest that ISVPs trigger caspase mediated apoptosis in MEFs and that WDR81 is required for this process.

**Figure 2.**
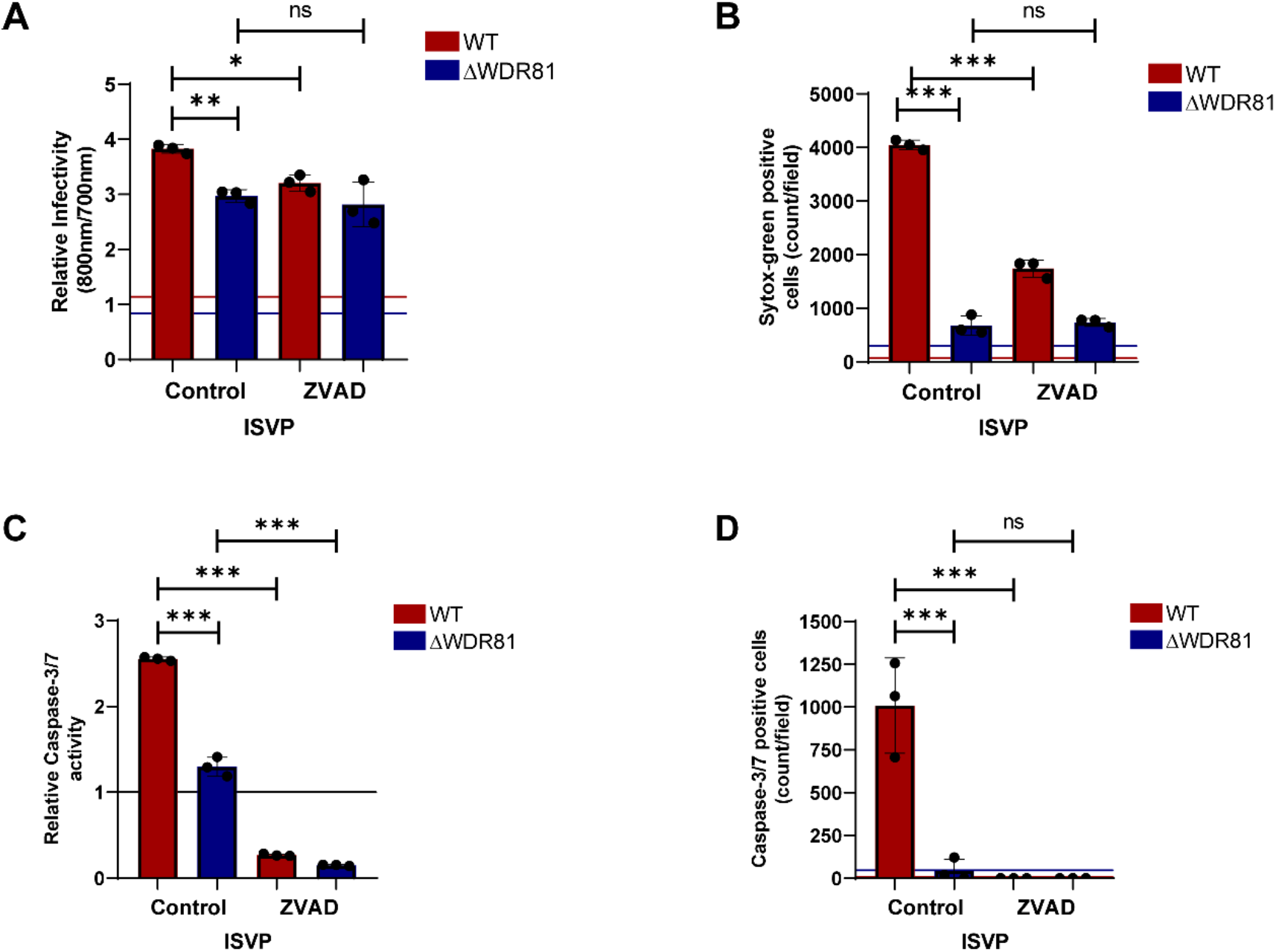
WDR81 regulates ISVP induced caspase-3/7 activation. (A) MEFs were adsorbed with reovirus T3D^CD^ ISVPs at MOI 10 PFU/cell. Cells were then incubated with either DMSO control or ZVAD-fmk (20µM) for 24 h at 37°C. Infectivity was measured in fixed cells by indirect immunofluorescence using LI-COR Odyssey Scanner. Relative infectivity was calculated using the ratio of signal at 800nm (reovirus) to signal at 700nm (cell control). Horizontal lines represent signals for mock infected cells. Error bars represent SD. **, P<0.005, *, P<0.05, by student t-test. (B) MEFs were adsorbed with reovirus T3D^CD^ ISVPs at MOI 10 PFU/cell. Cells were incubated for 48 h in media with either DMSO control or ZVAD-fmk (20µM) and with Sytox G (50 nM). Cell death was estimated 48 h post infection by analyzing the number of Sytox G positive cells per field. Horizontal lines represent signals for mock infected cells. Error bars represent SD. ***, P<0.0005 by student t-test. (C) MEFs were adsorbed with reovirus T3D^CD^ ISVPs at MOI 10 PFU/cell. Cells were incubated for 48 h in media with either DMSO control or ZVAD-fmk (20 µM). Caspase-3/7 activity was measured using Promega Caspase-3/7 Glo kit as per manufacturer’s instructions. Relative caspase-3/7 activity was calculated as the ratio of luminescence signal compared to background signal in mock infected cells. Horizontal line represents mock signal set to 1. Error bars represent SD. ***, P<0.0005 by student t-test. (D) MEFs were adsorbed with reovirus T3D^CD^ ISVPs at MOI 10 PFU/cell. Cells were incubated for 48 h in media with either DMSO control or ZVAD-fmk (20µM) and with Incucyte Caspase-3/7 dye (1:2000). The number of cells showing Caspase-3/7 activation per field was assessed by fluorescence microscopy. Horizontal lines represent signals for mock infected cells. Error bars represent SD. ***, P<0.0005 by student t-test.

### WDR81 deficient cells express genes that are also induced by TNF treatment

We hypothesized that the resistance of ΔWDR81 MEFs to ISVP-induced apoptosis might result from altered expression of genes regulating cell death and survival. Gene expression differences between WT and ΔWDR81 MEFs were compared by RNA-seq. RNA-seq analysis, using a false discovery rate (FDR) cutoff of <0.0001, revealed numerous genes differentially expressed between WT and ΔWDR81 MEFs (Fig. 3A). Ingenuity Pathway Analysis^28^ of differential gene expression revealed that WDR81 deficiency alters gene expression across multiple cellular pathways, including - but not limited to - those governing pathogen responses and cell-based immunity (data not shown). The large-scale, bidirectional changes in gene expression between WT and ΔWDR81 cells made it challenging to use these analyses to identify specific pathways driving the resistance of ΔWDR81 cells to ISVP-induced apoptosis. Many constitutively expressed host genes promote cell survival by repressing the activation of pathways of apoptosis^29^. One reason for the reduced ability of ΔWDR81 MEFs to undergo apoptosis following ISVP infection could be because such pro-survival genes are expressed at a higher level in these cells at a basal level. In MEFs, pro-survival genes can be upregulated by treatment with tumor necrosis factor (TNF)^29^. We therefore identified those genes whose expression is differentially regulated by TNF treatment (Fig 3B). We identified 107 TNF-responsive genes in WT cells (using the same stringent FDR cutoff). We then asked if these genes were also differentially expressed in ΔWDR81 MEFs. Notably, 66 of 107 TNF induced genes were differentially expressed at a basal level in untreated ΔWDR81 MEFs compared to untreated WT MEFs (Fig. 3C). To validate the differential gene expression analysis, transcript levels of A20 and Bcl2 – two TNF induced genes were assessed by RT-qPCR^30,31^. RT-qPCR analysis confirmed that ΔWDR81 MEFs exhibit ∼10-fold higher A20 mRNA levels and ∼2.5-fold higher Bcl2 mRNA levels compared to WT MEFs (Fig. 3D and 3E). These findings indicate that ΔWDR81 MEFs exhibit elevated basal expression of genes typically induced by TNF.

**Figure 3.**
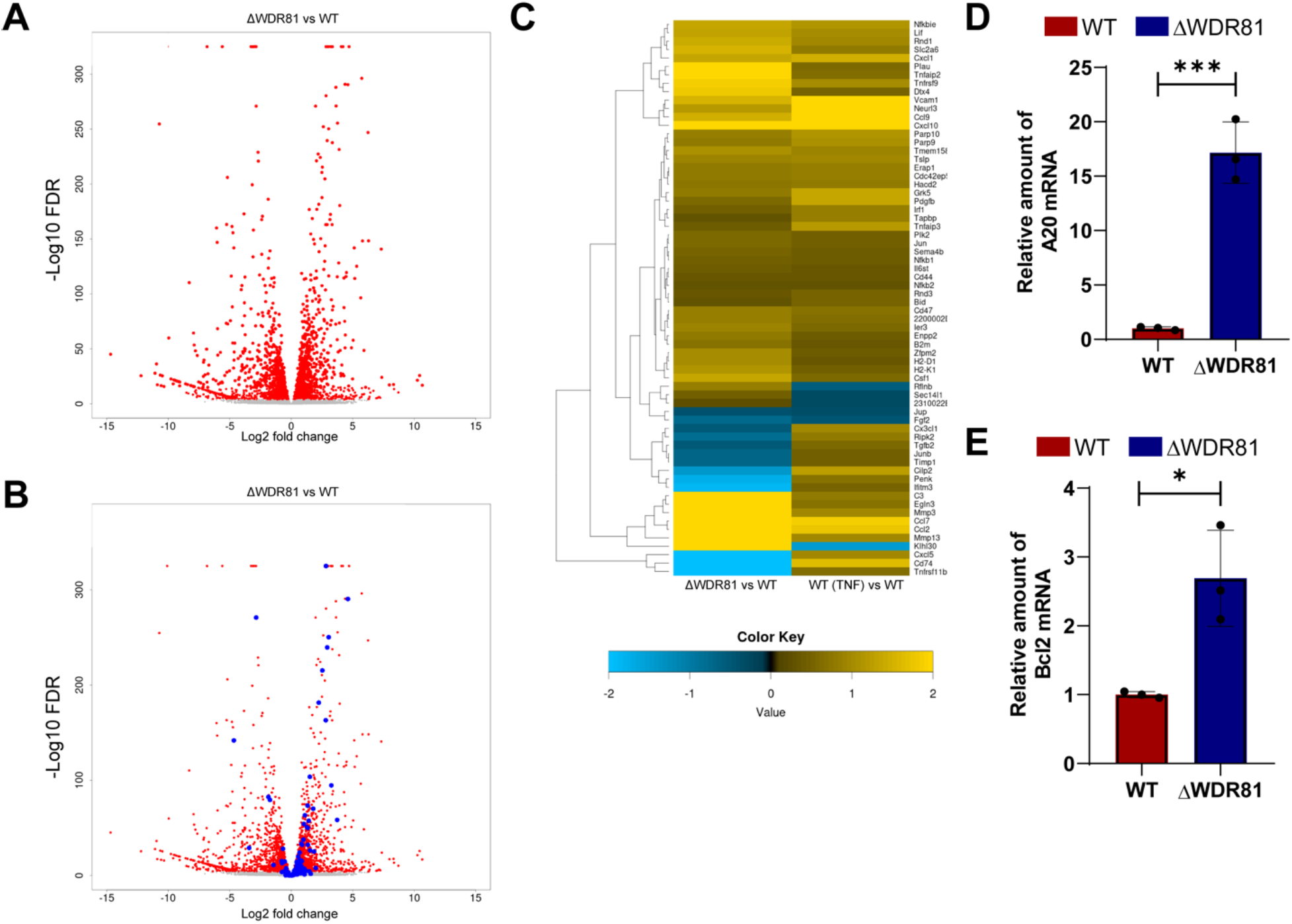
In the absence of WDR81, pro-survival gene expression is upregulated. (A) Total RNA was harvested from MEFs and gene expression was analyzed by RNA-seq. A volcano plot of genes differentially expressed in WT and ΔWDR81 is shown. Genes that are differentially expressed with FDR < 0.0001 are shown in red. (B) Genes in the volcano plot shown in A that are also differentially expressed in WT and TNF treated WT cells are marked in blue. (C) Heat map of 66 genes that are differentially expressed both in ΔWDR81 and WT MEFs as well as in TNF treated and control WT cells are shown. A scale bar showing log2 fold change in gene expression is included. (D) MEFs were grown in a 24 well plate and total mRNA was extracted using Bio-Rad Aurum Total RNA mini kit according to manufacturer’s protocol. cDNA was synthesized using Applied Biosystems High-Capacity cDNA Reverse Transcription kit. qPCR for A20 gene expression was done using Applied Biosystems StepOnePlus Real Time PCR system. A20 gene expression was calculated by 2^-ΔΔCT^ method using GAPDH as internal control. Error bars represent SD. ***, P<0.0005 by student t-test. (E) MEFs were grown in a 24 well plate and total mRNA was extracted using Bio-Rad Aurum Total RNA mini kit according to manufacturer’s protocol. cDNA was synthesized using Applied Biosystems High-Capacity cDNA Reverse Transcription kit. qPCR for Bcl2 gene expression was done using Applied Biosystems StepOnePlus Real Time PCR system. Bcl2 gene expression was calculated by 2^-ΔΔCT^ method using GAPDH as internal control. Error bars represent SD. *, P<0.05 by student t-test.

### Pro-survival gene expression in ΔWDR81 MEFs depends on IKK activity

After engaging its receptor on the cell surface, TNF signals via the IKK complex to activate NFκB and drive expression of its genomic targets^29^. Given the similarity between TNF-induced genes and those upregulated in ΔWDR81 MEFs, we hypothesized that IKK signaling may regulate the pro-survival gene expression observed in WDR81-deficient cells. Treatment with an IKK inhibitor significantly reduced A20 mRNA levels in ΔWDR81 MEFs (Fig. 4A). These data suggest that at least some of the genes that are differentially expressed in absence of WDR81 are controlled by IKK signaling. WDR81 negatively regulates endosomal type III PI3K, which generates phosphatidylinositol 3-phosphates (PtdIns3P) involved in endosomal signaling^24^. Consequently, ΔWDR81 cells are expected to exhibit elevated PI3K activity. To test if higher PI3K activity plays a role in higher pro-survival gene expression, WT and ΔWDR81 MEFs were treated with broad spectrum PI3K inhibitor LY294002^32^. Treatment of ΔWDR81 MEFs with LY294002 did not reduce A20 mRNA levels (Fig. 4B), indicating that elevated PI3K activity does not contribute to increased pro-survival gene expression in the absence of WDR81. These findings indicate that IKK signaling, but not PI3K activity, drives the upregulation of certain pro-survival genes in WDR81-deficient cells.

**Figure 4:**
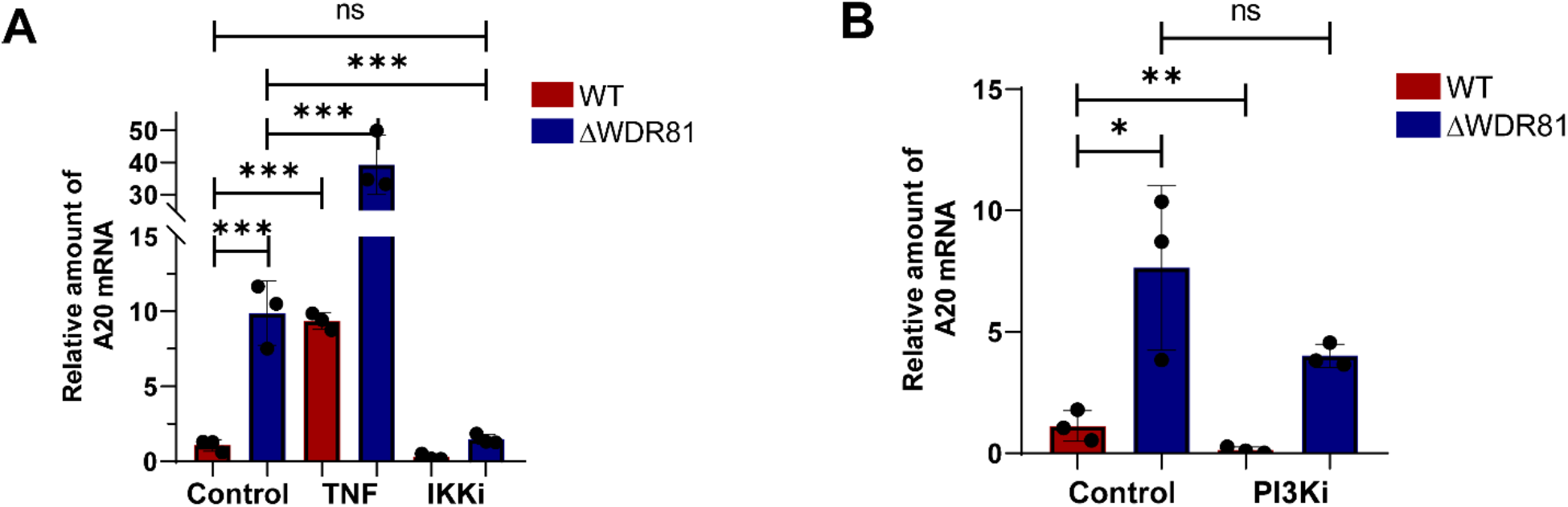
Higher A20 mRNA levels in WDR81 deficient cells are driven by IKK but not PI3K signaling. (A) MEFs were incubated in media with either DMSO control, TNF (50 ng/ml) or IKK inhibitor (5 μM) for three h. Total mRNA was extracted using Bio-Rad Aurum Total RNA mini kit according to manufacturer’s protocol. cDNA was synthesized using Applied Biosystems High-Capacity cDNA Reverse Transcription kit. qPCR for A20 gene expression was done using Applied Biosystems StepOnePlus Real Time PCR system. A20 gene expression was calculated by 2^-ΔΔCT^ method using GAPDH as internal control. Error bars represent SD. ***, P<0.0005 by one way ANOVA. (B) MEFs were incubated in media with DMSO control or PI3K inhibitor (25 μM) for three h. Total mRNA was extracted using Bio-Rad Aurum Total RNA mini kit according to manufacturer’s protocol. cDNA was synthesized using Applied Biosystems High-Capacity cDNA Reverse Transcription kit. qPCR for A20 gene expression was done using Applied Biosystems StepOnePlus Real Time PCR system. A20 gene expression was calculated by 2^-ΔΔCT^ method using GAPDH as internal control. Error bars represent SD. **, P<0.005, *, P<0.05 by one way ANOVA.

### WDR81 controls the activation of the extrinsic but not the intrinsic apoptotic pathway

TNF signaling via IKKs drives the expression of NFκB-dependent genes that enhance cell survival^29^. We hypothesized that the upregulated pro-survival genes in ΔWDR81 cells may confer resistance to apoptosis induced by other triggers. ABT-737 inhibits the Bcl2 family of antiapoptotic proteins^33^. Since the Bcl-2 family of proteins typically inhibits the activation of intrinsic apoptosis, ABT-737 treatment results in mitochondrial cytochrome c release to elicit the intrinsic apoptosis pathway. To test if WDR81 is required for apoptosis via the intrinsic pathway, we treated WT and ΔWDR81 MEFs with ABT-737 for 24 h. Treatment with ABT-737, resulted in comparable levels of cell death (Fig. 5A) and caspase-3/7 activation (Fig. 5B) in both WT and ΔWDR81 MEFs. These results indicate that WDR81 is dispensable for apoptosis induced by the intrinsic pathway. We next tested if WDR81 is required for extrinsic trigger induced apoptosis. To test this, we used TNF and cycloheximide (CHX) to induce death in WT and ΔWDR81 MEFs. This combination is a well characterized inducer of death-receptor mediated, extrinsic apoptosis^34^. While neither TNF, nor CHX alone altered cell survival (data not shown), upon treatment with the combination for 24 h, WT cells succumbed to cell death. In contrast, ΔWDR81 MEFs showed significantly lower number of dead cells (Fig. 5C). Similarly, TNF-CHX treatment was able to induce caspase-3/7 activity in WT MEFs but was unable to do the same in ΔWDR81 MEFs (Fig. 5D). Thus, WDR81 is required for apoptosis induced by extrinsic triggers. Together these data suggest that WDR81 is required for apoptosis induced by extrinsic but not intrinsic triggers.

**Figure 5:**
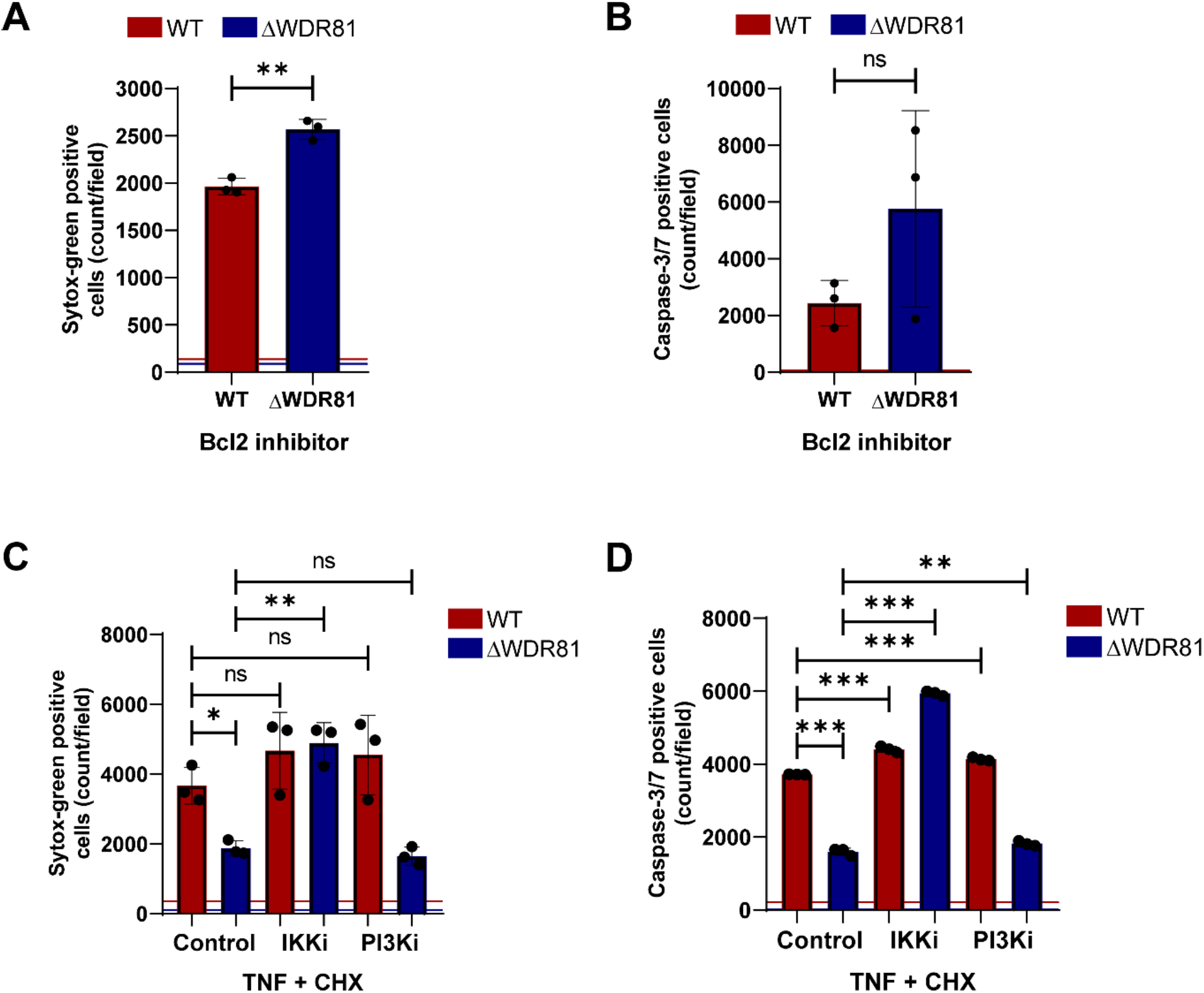
WDR81 is required for apoptosis activation by the extrinsic but not the intrinsic pathway. (A) MEFs were incubated in media with DMSO control or Bcl2 inhibitor (5 μM) and Sytox green (Sytox G)(50 nM) for 24 h. Cell death was quantified by analyzing the number of Sytox G positive cells per field. Horizontal lines represent signals for control DMSO treated cells. Error bars represent SD. **, P<0.005. (B) MEFs were incubated in media with DMSO control or Bcl2 inhibitor (5 μM) and Incucyte Caspase3/7 dye (1:2000) for 24 h. The number of cells showing Caspase-3/7 activation per field was assessed by fluorescence microscopy. Horizontal lines represent signals for control DMSO treated cells. Error bars represent SD. (C) MEFs were incubated in media supplemented with TNF (50 ng/ml) + CHX (10 μg/ml) and either DMSO control, IKK inhibitor (5 μM) or PI3K inhibitor (25 μM) for 24 h. Cell death was quantified by analyzing the number of Sytox G positive cells per field. Horizontal lines represent signals for control DMSO treated cells. Error bars represent SD. **, P<0.005, *, P<0.05 by one way ANOVA D: MEFs were incubated in media supplemented with Incucyte Caspase3/7 dye (1:2000) and TNF (50 ng/ml) + CHX (10μg/ml) and either DMSO control, IKK inhibitor (5μM) or PI3K inhibitor (25 μM) for 24 h. The number of cells showing Caspase-3/7 activation per field was assessed by fluorescence microscopy. Horizontal lines represent signals for control DMSO treated cells. Error bars represent SD. ***, P<0.0005 by one way ANOVA.

Since IKK and NFκB drive expression of pro-survival genes in ΔWDR81 cells, we assessed the impact of IKK inhibition on TNF-CHX induced cell death. We found that treatment with the IKK inhibitor restored TNF-CHX-induced cell death and caspase-3/7 activation in ΔWDR81 MEFs to levels comparable to WT MEFs (Fig. 5C, 5D). Thus, the resistance to extrinsic agonist triggered apoptosis in ΔWDR81 cells is driven by elevated IKK activity. Treatment with the PI3K inhibitor LY294002 to reduce elevated PI3K signaling in ΔWDR81 cells, did not restore TNF-CHX-induced cell death or caspase-3/7 activation in ΔWDR81 MEFs (Fig. 5C and 5D), even under conditions where the inhibitor was confirmed to be effective (data not shown). These results indicate that elevated PI3K activity does not contribute to the resistance of ΔWDR81 MEFs to extrinsic apoptosis. Furthermore, our results suggest that the elevated PI3K activity in the absence of WDR81 does not appear to impact IKK activity.

### WDR81 deficient cells support ISVP induced apoptosis when IKK activity is blocked

Apoptosis induction by reovirus requires death receptor signaling via TRAIL^17^ Additionally, for efficient cell death induction in MEFs, the death signals need to be amplified by Bid mediated activation of the intrinsic apoptotic pathway^15^. Given that IKK inhibition restores cell death in TNF-CHX treated cells, we asked if it would also restore cell death in ISVP infected cells. For these experiments, WT and ΔWDR81 MEFs were infected with reovirus ISVPs and subsequently incubated in the presence of IKK inhibitor. Infectivity measurements using indirect immunofluorescence revealed no significant differences in infection levels between DMSO-treated and IKK inhibitor-treated cells (Fig. 6A), confirming that the IKK inhibitor does impact viral replication. We measured cell death and caspase- 3/7 activation by ISVPs 48 h post infection in the presence of an IKK inhibitor. In control, DMSO-treated cells, ISVP infection induced a substantial level of apoptotic death in WT MEFs, while ΔWDR81 MEFs showed minimal cell death, consistent with our earlier observations (Fig. 6B). Although IKK inhibition had no effect on ISVP-induced apoptosis in WT MEFs, it fully restored apoptosis in ΔWDR81 MEFs to levels comparable to WT MEFs (Fig. 6B). Similarly, in the presence of IKK inhibitor, ISVPs induced caspase-3/7 activation in an equivalent number of cells to WT MEFs (Fig. 6C). Together, these results demonstrate that IKK driven pro-survival gene expression in ΔWDR81 MEFs prevents ISVP-induced caspase-3/7 activation and apoptosis.

**Figure 6:**
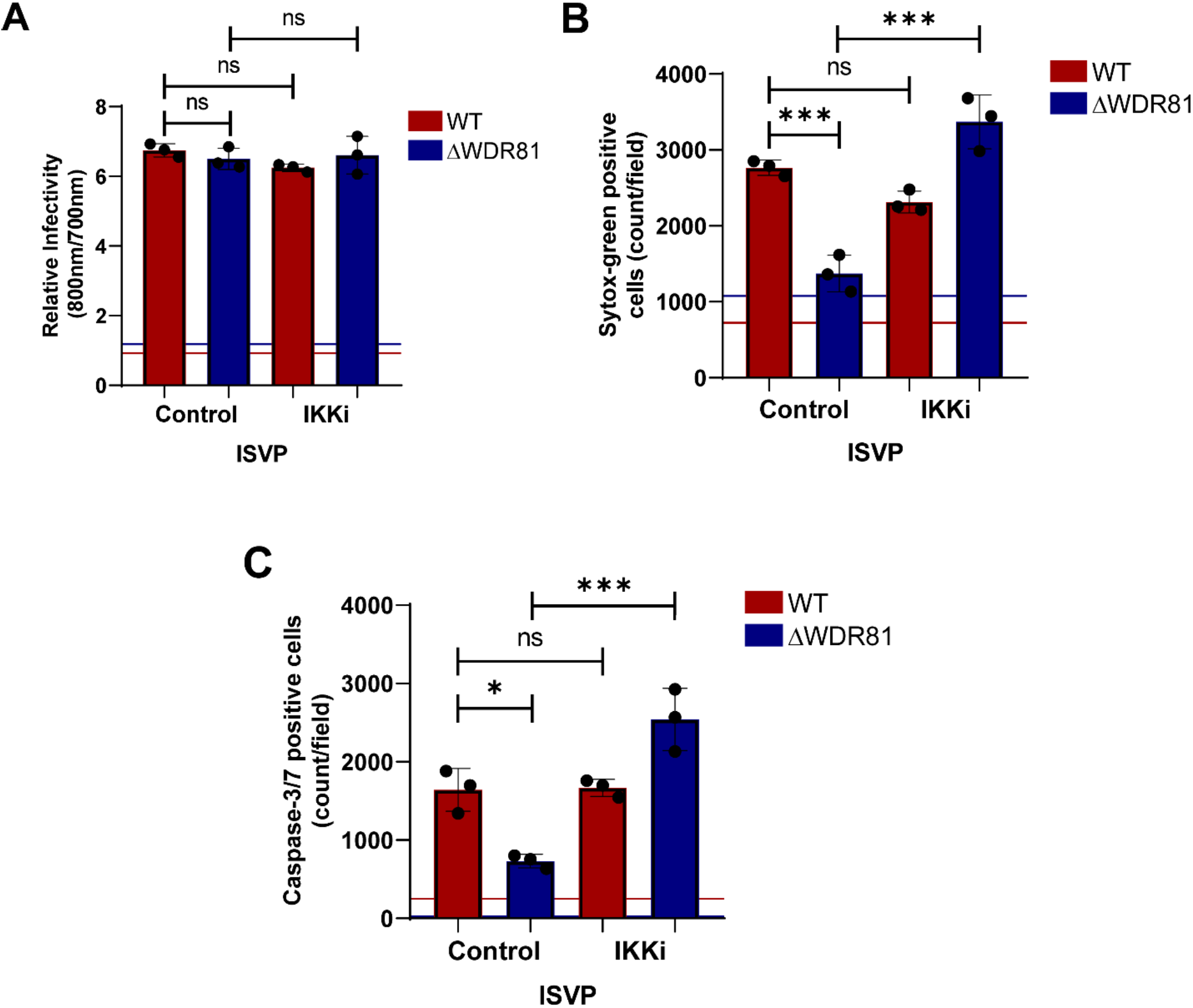
Resistance to ISVP induced cell death in ΔWDR81 MEFs can be overcome by blocking IKK activity. (A) MEFs were adsorbed with reovirus T3D^CD^ ISVPs at MOI 10 PFU/cell. Cells were then incubated for 24 h at 37°C in the presence of DMSO control or IKK inhibitor (5μM). Infectivity was measured in fixed cells by indirect immunofluorescence using LI-COR Odyssey Scanner. Relative infectivity was calculated using the ratio of signal at 800nm (reovirus) to signal at 700nm (cell). Horizontal lines represent signals for mock infected cells. Error bars represent SD. (B) MEFs were adsorbed with reovirus T3D^CD^ ISVPs at MOI 10 PFU/cell. Cells were then incubated for 48 h at 37°C in the presence of Sytox green (Sytox G) (50 nM) and either DMSO control or IKK inhibitor (5 μM). Cell death was quantified by analyzing the number of Sytox green positive cells per field. Horizontal lines represent signals for mock infected cells. Error bars represent SD. ***, P<0.0005 by one way ANOVA. (C) MEFs were adsorbed with reovirus T3D^CD^ ISVPs at MOI 10 PFU/cell. Cells were then incubated for 48 h at 37°C in the presence of Incucyte Caspase3/7 dye (1:2000) and either DMSO control or IKK inhibitor (5 μM). The number of cells showing Caspase-3/7 activation per field was assessed by fluorescence microscopy. Horizontal lines represent signals for mock infected cells. Error bars represent SD. ***, P<0.0005 by one way ANOVA.

## Discussion

In this study, we uncover a novel role of the protein WDR81 in regulating the extrinsic apoptosis pathway in cell culture. We show that WDR81 is required for the induction of apoptosis. In the absence of WDR81, cells infected with reovirus or those treated with TNF-CHX, a well characterized inducer of cell death via the extrinsic apoptotic pathway fail to activate caspase-3/7 and succumb to cell death (Fig. 1C and 5C). This role of WDR81 is specific to the extrinsic and not the intrinsic cell death pathway (Fig. 5A). In the absence of WDR81, although caspase-3/7 can be activated (Fig. 5B), their activation is impaired due to the upregulation of pro-survival signals (Fig. 4A and 5C). Our data indicate that a large majority of these upregulated pro-survival signals are those that are regulated by the IKK-NFκB pathway (Fig. 3B, 3C, 4A, 5C and 6A). Inhibition of the IKK pathway is sufficient to allow extrinsic triggers like reovirus or TNF-CHX to mediate caspase-3/7 activation and death (Fig. 5C, 5D, 6A and 6B). Based on these data, we propose a model (Fig. 7) where WDR81 is responsible for the basal suppression of IKK-NFκB pro-survival genes. Our data suggests that in the absence of WDR81, upregulated pro-survival genes prevent reovirus ISVPs and TNF from activating apoptosis. This work highlights a novel association between WDR81 and the IKK-NFκB mediated cell survival pathway.

**Figure 7.**
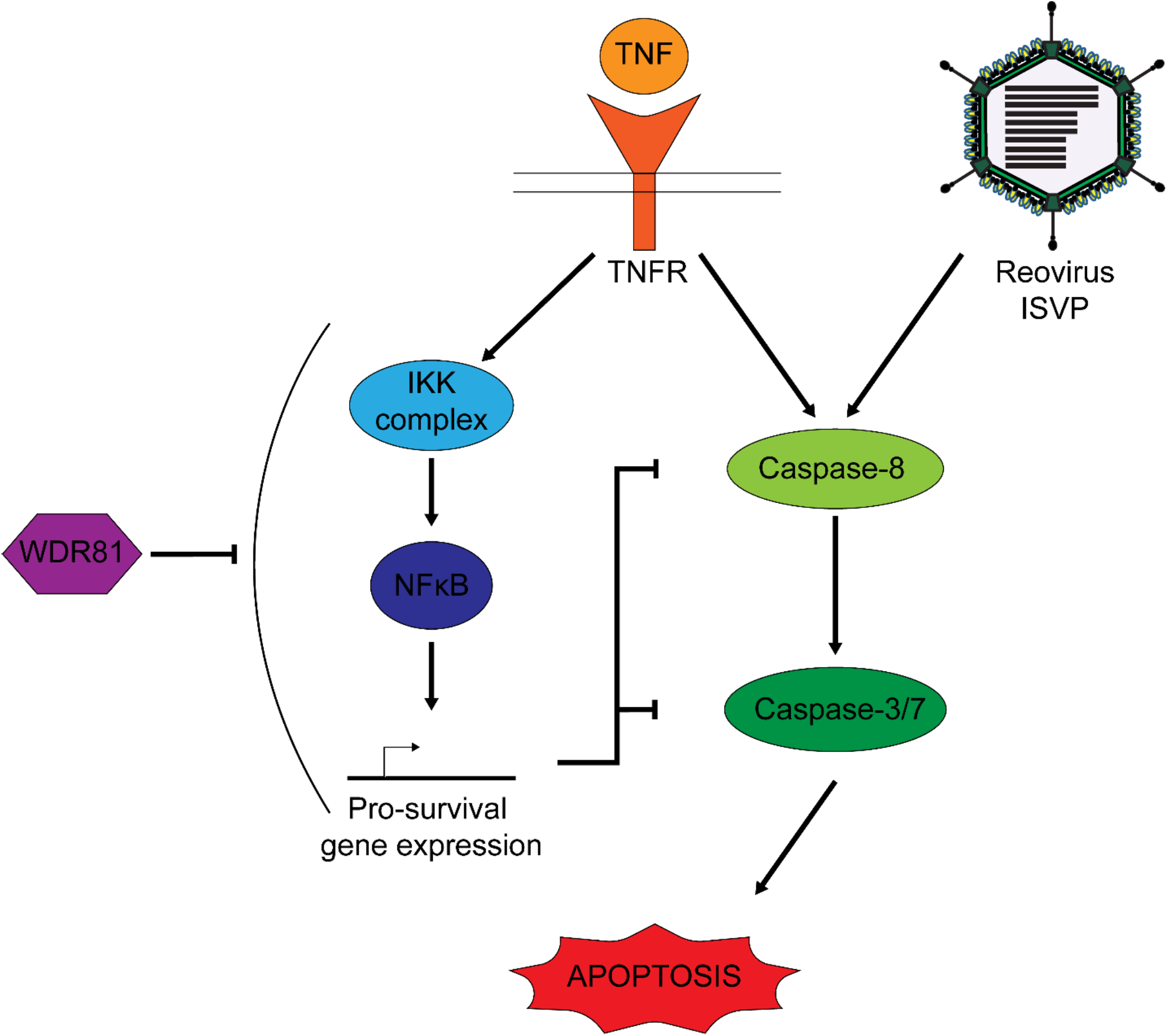
Proposed model for the role of WDR81 in reovirus and TNF induced apoptosis. Infection with reovirus ISVPs induces apoptosis through the activation of caspase-8 and caspase-3/7. TNF treatment upregulates pro-survival gene expression through IKK-NFκB pathway which inhibits its ability to trigger apoptosis. Treatment of cells with TNF+CHX (which inhibits translation of pro-survival genes, allows TNF to activate apoptosis. Our data support the model that WDR81 regulates cell death by suppressing IKK-NFκB mediated expression of pro-survival genes.

WDR81 is a cellular protein that inhibits Class III PI3Ks on the early endosomal membrane^24^. This function regulates the efficient conversion of early to late endosomes and thus cellular endosomal trafficking. Cells lacking WDR81 have delayed early to late endosome conversion and lysosomal turnover of plasma membrane proteins such as epidermal growth factor receptor is impaired^24^. Since PI3K inhibitors were not able to rescue cell death in ΔWDR81 MEFs, we think that the cell survival controlling function of WDR81 is independent of its role as a PI3K inhibitor. WDR81 can interact with p62 and LC3C promoting autophagy^35^. In the absence of WDR81, there is an increase in protein aggregation and reduced autophagic clearance. It is possible that in the absence of WDR81, there is decreased autophagic turnover of an activator of the IKK-NFκB pathway. Mutations in WDR81 have been associated with increased cell death in purkinje cells of mice, resulting in improper motor abilities^36^. This suggests that WDR81 may have opposite functions in different cells.

Previous work has shown that NFκB signaling mediated through the IKK complexes is required for reovirus induced death^37,38^. These data were obtained using cells deficient in IKK or NFκB subunits. Reovirus affects NFκB in two phases. In the first phase, infection triggers NFκB activation and in the second stage NFκB activity is inhibited^39^. It is thought that NFκB inhibition later in infection decreases the concentration of survival factors in the cell thereby sensitizes them to cell death^39^. Since WDR81-deficient cells have constitutively high IKK-NFκB activity, they allow us to explore the biological effect of not being able to effectively block NFκB later in infection. Our work demonstrating that WDR81-deficient cells are resistant to apoptosis supports the idea that the second stage inhibition of NFκB is required for cell death. It is possible NFκB inhibition later in infection occurs in a cell type specific manner. Such a possibility may account for organ specific role of NFκB in reovirus pathogenesis^40^.

Reovirus triggers death via TNF related apoptosis inducing ligand (TRAIL) mediated signaling^11,12^. In reovirus infected cells, TRAIL signaling promotes casapase-8 activation^17^. But caspase-8 activation is not sufficient to induce apoptosis. Apoptosis induction after reovirus requires casapse-8 mediated cleavage of Bid^15^. Cleaved Bid (tBid) engages the mitochondrial apoptotic pathway to induce sufficient effector caspase activation to induce cell death^41^. TNF-CHX shares some similarities with cell death signaling with reovirus since it also induces caspase-8 activation. Following TNF-CHX treatment, whether amplification of death signals via the intrinsic mitochondrial apoptotic pathway is needed for cell death may vary with cell type ^42^. Our results also indicate that WDR81-deficient cells are capable of undergoing cell death following direct activation of the intrinsic pathway through inhibition of Bcl-2 family of anti-apoptotic proteins. Thus, the effect of WDR81 on cell death appears to be restricted to events in death receptor signaling that lead to caspase-8 activation. IKK dependent NFκB mediated gene expression drives the expression of many proteins that are inhibit caspase-8 activation and promote cell survival^29^. Interestingly, one such target, the A20 protein is sufficient to protect cells from TNF induced cell death^29^. While our data demonstrate that A20 is expressed at a higher level in absence of WDR81, whether the enhanced expression of A20 alone is sufficient to prevent efficient cell death in WDR81-deficient cells remains unknown and a subject of our investigation.

## Materials and Methods

### Cells and Viruses

Murine L929 cells (spinner cells) were grown at 37°C in Joklik’s Minimal Essential Medium (Lonza) supplemented with 5% fetal bovine serum (FBS) (Life Technologies), 2mM L-Glutamine (Invitrogen), 0.5 U/ml penicillin, 50 μg/ml streptomycin (Sigma Aldrich), and 25 ng/ml amphotericin B (Sigma-Aldrich). Mouse embryo fibroblasts (MEFs) containing non-targeting plasmid (WT MEFs) and cells containing lentiCRIPSPRv2 vector targeting WDR81 (ΔWDR81 MEFs) were maintained at 37°C in Dulbecco’s Modified Eagle Medium (DMEM) (Gibco) supplemented with 10% FBS, 2mM L-Glutamine and 2 μg/ml puromycin (Invivogen). All reovirus experiments were performed with Type 3 Dearing from the Cashdollar laboratory (T3D^CD^).

### Reovirus Purification

Reovirus T3D^CD^ was propagated in spinner cells. Spinner cells infected with second passage of reovirus stock were lysed by sonication. Virions were extracted from lysates using Vertrel-XF specialty fluid (Dupont). The extracted particles were layered onto 1.2- to 1.4 g/cm^3^ CsCl step gradients. Gradients were centrifuged at 187, 000 x g for 4 h at 4°C. Bands corresponding to purified virions (1.36 g/cm^3^) were isolated and dialyzed into virus storage buffer (10 mM Tris [pH 7.4], 15 mM MgCl2, and 150 mM NaCl). After dialysis, the particle concentration was determined by measuring the optical density at 260 nm (OD260) of the purified virion stocks (1 unit at OD260 = 2.1 x 10^12^ particles/ml). The purification of virions was confirmed by SDS-PAGE^43^.

### Generation of reovirus ISVPs

Purified T3D^CD^ virions (2x10^11^ particles/ml) were digested with 200μg/ml Na-p-tosyl-L-lysine chloromethyl ketone (TLCK)-treated chymotrypsin (Worthington Biochemical) in a total volume of 100 μl for 30 min at 32°C. The reactions were then incubated on ice for 15 mins and quenched by adding 1mM phenylmethylsulphonyl fluoride (Sigma-Aldrich). ISVP generation was confirmed by SDS-PAGE^44^.

### Reovirus infectivity by indirect immunofluorescence

MEFs were seeded in 96 black well plates (Greiner) at 2x10^4^ cells/well. 24 h after seeding, cells were adsorbed with T3D^CD^ virions or ISVPs at MOI of 10 pfu/cell for 1 hour at room temperature (RT) using PBS for mock infections. Inoculum was then replaced with media and incubated at 37°C in CO_2_ incubator. 24 h post infection, cells were washed with PBS and fixed with 100 µl methanol for 30 minutes at -20°C. Cells were then washed with 0.5% PBS-Tween 20 (PBS-T) and blocked using 1% PBS-Bovine serum albumin (PBS-BSA). Subsequently cells were stained for 1 hour with reovirus polyclonal serum (1:1000 in 1% PBS-BSA). Cells were then washed with PBS-T three times before staining for 1 hour with LICOR donkey anti-rabbit IRDye-800 CW (1:1000 in 1% PBS-BSA) and Draq-5 (1:10,000 in 1% PBS-BSA). Cells were then washed three times with PBS-T and 50 µl water added each well. Plate was imaged using LICOR-Odyssey CLX scanner to capture IR images at 800 nm (reovirus) and 700 nm (cells). Infectivity was measured by signal at 800 nm (reovirus) normalized to signal at 700 nm (cells).

### Single step reovirus growth assay

MEFs were seeded in 24 well plates (Greiner) at 1x10^5^ cells/well. 24 h after seeding, cells were adsorbed with T3D^CD^ virions or ISVPs at MOI of 10 pfu/cell for 1 hour at room temperature (RT). Inoculum was then replaced with media as described above and incubated at 37°C in CO_2_ incubator. 24 h post infection, cells were harvested by 3x freeze-thaw cycle and virus titer was measured by plaque assay on spinner cells. Briefly, confluent 6 well plates of spinner cells were adsorbed with 100 µl of 10-fold serially diluted, freeze-thawed lysate for 1 hour. Cells were then overlayed with Media 199 supplemented with 2mM L-Glutamine (Invitrogen), 100 μg/ml streptomycin (Invitrogen), 25 ng/ml amphotericin B (Sigma-Aldrich) and 1% Difco Bacto Agar (BD). Plates were incubated until countable plaques were visible and fixed with 3.7% formaldehyde in PBS. Fixed plates were then stained with 1% Crystal Violet in 5% ethanol solution to visualize and count plaques.

### Cell death assays

MEFs were seeded in 96 black well plates (Greiner) at 2x10^4^ cells/well. 24 h after seeding, cells were adsorbed with T3D^CD^ virions or ISVPs at MOI of 10 pfu/cell for 1 hour at room temperature (RT) using PBS for mock infections. Inoculum was replaced with media as described above and supplemented with 50 nM Sytox-green nucleic acid dye (Invitrogen). 48 h post infection plates were imaged by fluorescence microscopy using Incucyte (Essen Biosciences). Sytox green positive cells per field were quantified using the associated software.

### Caspase-3/7 assays

For fluorescence microscopic analysis of caspase-3/7, MEFs were seeded in 96 black well plates (Greiner) at 2x10^4^ cells/well. 24 h after seeding, cells were adsorbed with T3D^CD^ virions or ISVPs at MOI of 10 pfu/cell for 1 hour at room temperature (RT) using PBS for mock infections. Inoculum was replaced with media as described above supplemented with 2.5 µM Incucyte Caspase-3/7 Green (Sartorius). 48 h post infection plates were imaged by fluorescence microscopy using Incucyte S3 imager (Essen Biosciences). Caspase-3/7 positive cells per field were quantified using the associated software.

For luminescence based relative caspase-3/7 assay, MEFs were seeded in 96 black well plates (Greiner) at 2x10^4^ cells/well. 24 h after seeding, cells were adsorbed with T3D^CD^ virions or ISVPs at MOI of 10 pfu/cell for 1 hour at room temperature (RT) using PBS for mock infections. Inoculum was replaced with media as described above. 48 h post infection, caspase-3/7 assay was measured using Promega Caspase-3/7 Glo kit according to manufacturer’s instructions. Luminescence was measured using Biotek SynergyLX multimode reader. Relative caspase-3/7 activity was calculated by normalizing luminescence signal of sample to that of mock, which was set to 1.

### Analysis of host gene expression by RT-qPCR

Total RNA was extracted from cells using Bio-Rad Aurum Total RNA minikit according to manufacturer’s instructions. Eluted RNA was converted into cDNA using Thermo-fisher High-Capacity cDNA Reverse transcription kit using random hexamers. A20 and Bcl2 gene expression was estimated by qPCR using Applied Biosystems Stepone Realtime PCR machine. Relative mRNA levels were calculated using GAPDH as internal control using 2^-ΔΔCT^ method^45^. Primers used for qPCR are as follows: GAPDH, For ACCCAGAAGACTGTGGATGG and Rev GGATGCAGGGATGATGTTCT, A20, For CTGGATGTCAATCAACAATGGGA and Rev ACTAGGGTGTGAGTGTTTTCTGT, Bcl2, For ATGCCTTTGTGGAACTATATGGC and Rev GGTATGCACCCAGAGTGATGC.

### Library preparation for RNA-seq and gene expression analysis

Total RNA was extracted from cells using Bio-Rad Aurum Total RNA minikit according to manufacturer’s instructions. RNA was submitted to Indiana University Center for Genomics and Bioinformatics for RNA-seq processing. Briefly, cDNA library construction was done using a TruSeq stranded mRNA low-throughput (LT) sample prep kit (Illumina) following manufacturers protocol. Sequencing was performed using an Illuniman NextSeq 500 platform with a 75-bp sequencing module generating 38-bp paired end reads. After the sequencing run demultiplexing was performed with bcl2fastq v2.20.0.422.

The sequenced reads were adapter trimmed, and quality filtered using Trimmomatic ver. 0.38^46^ setting the cutoff threshold for average base quality score at 20 over a window of 3 bases, with a minimum read length of 20 bases after trimming (parameters: LEADING:20 TRAILING:20 SLIDINGWINDOW:3:20 MINLEN:20). Cleaned reads were mapped to the mouse genome sequence GRCm39 using STAR RNA-seq aligner version 2.7.11a^47^ using default parameters. Concordantly mapped read pairs aligning to the exon regions of annotated genes on the sense strand (Ensembl release-109) were counted using featureCounts tool version 2.0.0^48^ of subread package (parameters: -t exon -g gene_id -s 2 - p -B -C). The differential expression analysis was conducted using DESeq2 version 1.44.0^49^.

### Chemicals and Reagents

Pan-caspase inhibitor ZVAD-fmk (Cayman chemicals) was used at a final concentration of 20 µM. IKK inhibitor BAY-65-1942 (Bayer) was used at a final concentration of 5 nM. PI3K inhibitor LY294002 (Cayman chemicals) was used at a final concentration of 25 nM. Human tissue necrotic factor alpha (TNF) (Sigma-Aldrich) was used at a final concentration of 50 ng/ml. Cycloheximide was used at a final concentration of 10 µg/ml. Bcl2 inhibitor ABT-737 (Cayman chemicals) was used at a final concentration of 5 μM.

## Notes

### Competing Interest Statement

The authors have declared no competing interest.

